# Mapping the corticoreticular pathway from cortex-wide anterograde axonal tracing in the mouse

**DOI:** 10.1101/2021.06.23.449661

**Authors:** Pierce Boyne, Oluwole O. Awosika, Yu Luo

**Author notes:** **Corresponding author:** Pierce Boyne, PT, DPT, PhD, NCS, Health Sciences Building, room 267, 3225 Eden Ave. Cincinnati, OH, 45267-0394.

## Abstract

The corticoreticular pathway (CRP) has been implicated as an important mediator of motor recovery and rehabilitation after central nervous system damage. However, its origins, trajectory and laterality are not well understood. This study mapped the mouse CRP in comparison with the corticospinal tract (CST). We systematically searched the Allen Mouse Brain Connectivity Atlas (© 2011 Allen Institute for Brain Science) for experiments that used anterograde tracer injections into the right isocortex in mice. For each eligible experiment (N=607), CRP and CST projection strength were quantified by the tracer volume reaching the reticular formation motor nuclei (RF_motor_) and pyramids respectively. Tracer density in each brain voxel was also correlated with RF_motor_ versus pyramids projection strength to explore the relative trajectories of the CRP and CST. We found significant CRP projections originating from the primary and secondary motor cortices, anterior cingulate, primary somatosensory cortex and medial prefrontal cortex. Compared with the CST, the CRP had stronger projections from each region except the primary somatosensory cortex. Ipsilateral projections were stronger than contralateral for both tracts (above the pyramidal decussation), but the CRP projected more bilaterally than the CST. The estimated CRP trajectory was anteromedial to the CST in the internal capsule and dorsal to the CST in the brainstem. Our findings reveal a widespread distribution of CRP origins and confirm strong bilateral CRP projections, theoretically increasing the potential for partial sparing after brain lesions and contralesional compensation after unilateral injury.

**SIGNIFICANCE:** The corticoreticular pathway (CRP) provides volitional input to brainstem nuclei that generate walking command signals, facilitate balance and direct limb movements. Upregulation of this pathway appears to be a central mechanism of movement recovery after brain and spinal cord injury, but its anatomy is not well understood. We showed that the mouse CRP originates from widespread parts of the cortex, including non-motor regions, that it projects strongly to both sides of the brainstem, and that its projections are more distributed and bilateral than the corticospinal tract. These findings suggest that the CRP may be particularly resilient to complete disruption.

## INTRODUCTION

The reticulospinal tract is a major motor pathway that delivers the primary input to locomotion-generating circuits in the spinal cord, while also supporting anticipatory postural control and efficient motor synergies in the upper and lower limbs.(Brownstone and Chopek, 2018; Matsuyama et al., 2004; Riddle et al., 2009) The corticoreticular pathway (CRP) provides volitional input to the reticulospinal system via direct and indirect projections from the cerebral cortex to the medial reticular formation motor nuclei (RF_motor_).(Fisher et al., 2021; Jinnai, 1984; Matsuyama et al., 2004) While the CRP has received much less attention than the more recently evolved corticospinal tract (CST),^e.g.^(Capaday et al., 1999; Dawes et al., 2008; Jayaram et al., 2012; Stinear et al., 2007) accumulating evidence in animal studies (including non-human primates) now indicates that the CRP is likely a critical mediator of motor recovery after central nervous system damage, that also mediates effects of rehabilitation.(Asboth et al., 2018; Darling et al., 2018; Glover and Baker, 2020; Ishida et al., 2019; Takase et al., 2017; Zaaimi et al., 2012; Zaaimi et al., 2018) However, the origins, trajectory and laterality of the CRP are not well understood.

In humans, white matter tracts like the CRP are typically mapped by simulating streamlines through pre-processed diffusion-weighted magnetic resonance images. While this method provides useful non-invasive, *in vivo* measures, it is limited by insufficient resolution to resolve axonal bundles and inability to determine the direction of neural conduction.(Calamante, 2019) Consequently, it is crucial for diffusion tractography to be guided by anatomical knowledge in order to maximize tract coverage and minimize inclusion of false pathways.(Aydogan et al., 2018; Azadbakht et al., 2015; Gutierrez et al., 2020) Unfortunately, this anatomical knowledge is incomplete for the CRP, which limits confidence in prior human CRP mapping.

Most studies in humans and other animals have exclusively mapped CRP projections from the primary and secondary motor areas.(Asboth et al., 2018; Darling et al., 2018; Fisher et al., 2021; Fregosi et al., 2017; He and Wu, 1985; Ishida et al., 2019; Jang and Lee, 2019; Jinnai, 1984; Kably and Drew, 1998a; Kably and Drew, 1998b; Lamas et al., 1994; Matsuyama et al., 2004; Pilyavsky, 1975; Schulz et al., 2017; Takase et al., 2017; Yeo et al., 2020) However, the extent of cortical inputs to the CRP has not been well defined, and there have been preliminary indications of potentially important CRP bundles originating from outside the motor cortex.(Keizer et al., 1989; Newman et al., 1989; Rho et al., 1997; Rossi and Brodal, 1956; Shammah-Lagnado et al., 1987) For example, one anterograde axonal tracing study using wheat germ agglutinin-horseradish peroxidase (WGA-HRP) in the rat revealed dense RF_motor_ projections from the anterior cingulate cortex (N=2) and medial prefrontal cortex (N=5).(Newman et al., 1989) Likewise, a classical retrograde tracing experiment using HRP or WGA-HRP injections into RF_motor_ in rats (N=41) found that the strongest CRP projections originated from the anterior cingulate and medial prefrontal cortices.(Shammah-Lagnado et al., 1987) Such projections could be important because they may provide greater opportunity for CRP sparing after a brain lesion, and novel targets for neuromodulation. However, other small studies (N=2-14) using anterograde lesion degeneration mapping or classical retrograde tracer injections into the medial reticular formation in the cat or primate have only observed *either* anterior cingulate(Keizer et al., 1989) *or* medial prefrontal(Rho et al., 1997; Rossi and Brodal, 1956) CRP origins, or neither.(Berrevoets and Kuypers, 1975; Keizer et al., 1984)

Uncertainty about the extent of CRP origins has also been compounded by the limitations of these prior tract tracing experiments testing the breadth of its inputs, which were all from the 20^th^ century. Classical tracers commonly have spread from the injection site, non-specific cell labelling, bidirectional transport (anterograde and retrograde) and trans-synaptic travel, which could have led to false positives.(Saleeba et al., 2019) Likewise, false negatives may have occurred due to insufficient neuronal uptake or transport over long-range CRP axons,(Saleeba et al., 2019) a limited search window or a differing taxonomy of brain regions. Modern tract tracing methods are largely able to overcome these limitations,(Chamberlin et al., 1998; Oh et al., 2014; Ragan et al., 2012; Saleeba et al., 2019; Wang et al., 2020) but the extent of cortical inputs to the CRP has not been previously assessed systematically with next-generation viral tracers or localization procedures.

In addition, the laterality of the CRP also remains incompletely understood. The CRP has been found to project bilaterally to RF_motor_,^e.g.^(Fisher et al., 2021; Fregosi et al., 2017; Kably and Drew, 1998; Matsuyama and Drew, 1997; Rho et al., 1997) which could have important implications for recovery from unilateral brain lesions.(Brownstone and Chopek, 2018; Jang and Lee, 2019; Takase et al., 2017) However, a paucity of studies have quantified the laterality of these projections,(Fisher et al., 2021; Kably and Drew, 1998; Rho et al., 1997) and those studies have only tested small numbers of neurons originating from restricted sites within the motor cortex,(Fisher et al., 2021; Kably and Drew, 1998) and/or have had relatively small sample sizes (N=4-14).(Fisher et al., 2021; Kably and Drew, 1998; Rho et al., 1997) A broader quantification of CRP lateralization would provide a better understanding of the proportion of contralateral projections available for compensation after unilateral brain injury.

The Allen Mouse Brain Connectivity Atlas(Oh et al., 2014) (© 2011 Allen Institute for Brain Science http://connectivity.brain-map.org/) provides a unique opportunity to address these knowledge gaps. This open data resource was generated from brain-wide projection mapping, using stereotaxic injections with enhanced green fluorescent protein (EGFP)-expressing adeno-associated viral vectors (rAAV2/1) for anterograde axonal labelling, with negligible retrograde transport.(Chamberlin et al., 1998) Serial two-photon tomography(Ragan et al., 2012) was then followed by spatial registration into a common three-dimensional reference space (CCFv3).(Wang et al., 2020) Labeled pixel volumes were quantified at the injection site and in bilateral atlas regions across the brain, including many specific reticular nuclei.(Oh et al., 2014) This resource has already made major contributions to our understanding of brain organization,(Oh et al., 2014) and it currently includes data from 2,994 anterograde tracer experiments, but it has not been previously used to evaluate the CRP. The current study leveraged the Allen Connectivity Atlas to determine the location, extent and laterality of cortical inputs to the CRP in the mouse, while also mapping the cortical inputs to the CST for comparison. In addition, we used statistical methods developed for human brain imaging analysis to explore the trajectory of the CRP relative to the CST.

## METHODS AND MATERIALS

### Data sources

Anterograde tracer data were obtained from the Allen Mouse Brain Connectivity Atlas(Oh et al., 2014) (© 2011 Allen Institute for Brain Science http://connectivity.brain-map.org/). The experimental procedures that produced this dataset are summarized in the introduction above and were approved by the Institutional Animal Care and Use Committee of the Allen Institute for Brain Science.(Oh et al., 2014) Experiments were performed on adult mice with injections at postnatal day 54-58 and euthanasia 21 days later followed by brain imaging and data processing.(Oh et al., 2014) Further details on the experimental procedures have been described in previous publication.(Oh et al., 2014) We also obtained reference anatomical data and brain region annotation labels from the Allen Mouse Brain Atlas(Sunkin et al., 2006) (© 2004 Allen Institute for Brain Science http://atlas.brain-map.org/) in the CCFv3 reference space, from the Scalable Brain Atlas(Bakker et al., 2015) (https://scalablebrainatlas.incf.org/mouse/ABA_v3).

### Data selection & processing

To identify all potentially eligible experiments in the connectivity atlas, we performed a ‘source search’ for those that included injection into the cerebral isocortex using the anterograde EGFP tracer. The following link reproduces the search: http://connectivity.brain-map.org/projection?searchMode=source&sourceDomain&=315&primaryStructureOnly=true&tracers=10&isi=false. A ‘target search’ within those results was then performed to find experiments where tracer was detected in one or more of the reticulospinal motor nuclei involved with limb movement (RF_motor_; Table 1; http://connectivity.brain-map.org/projection?searchMode=target&sourceDomain=315&primaryStructureOnly=true&tracers=10&isi=false&targetDomain=1048,307,938,970,978,852,395,1098,1107,136,1093,146,880,162,358,599626927&hemisphere=either&targetVolumeThreshold=0.0000).(Brownstone and Chopek, 2018)

**Table 1.**
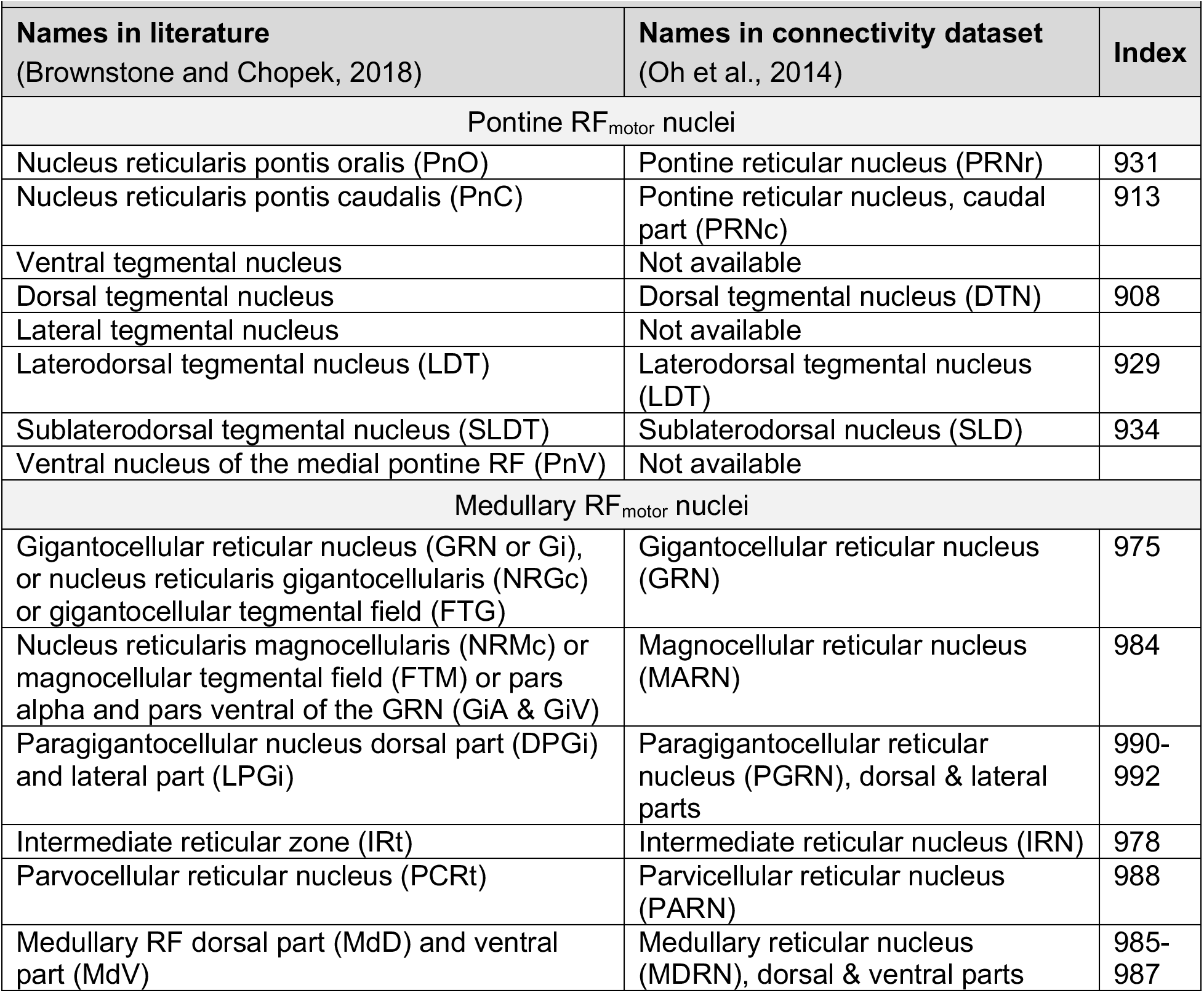
Reticular formation motor nuclei involved with limb movement (RF_motor_) which form the origin of the reticulospinal tracts.(Brownstone and Chopek, 2018)

One experiment (ID: 249396394) was excluded because the injection site mapped to the superior colliculus (outside of the isocortex). We also restricted our analysis to injection experiments in the right hemisphere, because there was minimal coverage of non-visual regions in the left hemisphere. Since cortical layer 5 is the origin of corticofugal projection pathways like the CRP and CST,(Gerfen et al., 2018) we also excluded experiments that used transgenic mice with limited expression in layer 5, according to the documentation (http://connectivity.brain-map.org/transgenic). This meant that experiments using the following transgenic lines were excluded: Calb1-T2A-dgCre, Calb2-IRES-Cre, Cort-T2A-Cre, Crh-IRES-Cre_BL, Ctgf-T2A-dgCre, Cux2-CreERT2, Cux2-IRES-Cre, Erbb4-T2A-CreERT2, Esr1-2A-Cre, Gad2-IRES-Cre, Grp-Cre_KH288, Htr3a-Cre_NO152, Nos1-CreERT2, Nr5a1-Cre, Ntsr1-Cre_GN220, Oxtr-T2A-Cre, Penk-IRES2-Cre-neo, Pvalb-IRES-Cre, Slc18a2-Cre_OZ14, Syt17-Cre_NO14 and Tac1-IRES2-Cre.

Included experiments used wild-type mice (C57BL/6J) or one of the following transgenic lines: A930038C07Rik-Tg1-Cre, Adcyap1-2A-Cre, Cart-Tg1-Cre, Chrna2-Cre_OE25, Chrnb4-Cre_OL57, Drd3-Cre_KI196, Efr3a-Cre_NO108, Etv1-CreERT2, Glt25d2-Cre_NF107, Gnb4-IRES2-Cre, Gnb4-IRES2-CreERT2, Gng7-Cre_KH71, Gpr26-Cre_KO250, Grm2-Cre_MR90, Htr2a-Cre_KM207, Npr3-IRES2-Cre, Ntng2-IRES2-Cre, Oxtr-Cre_ON66, Plxnd1-Cre_OG1, Pvalb-T2A-CreERT2, Rasgrf2-T2A-dCre, Rbp4-Cre_KL100, Rorb-IRES2-Cre, Rorb-IRES2-Cre-neo, Scnn1a-Tg3-Cre, Sepw1-Cre_NP39, Sim1-Cre_KJ18, Slc17a8-iCre, Slc17a8-IRES2-Cre, Slc32a1-IRES-Cre, Sst-IRES-Cre, Syt6-Cre_KI148, Tlx3-Cre_PL56, Trib2-F2A-CreERT2, Vip-IRES-Cre.

The projection strength from each cortical injection site to RF_motor_ was calculated by dividing the total volume of tracer-labelled pixels in RF_motor_ by the injection site volume.(Oh et al., 2014) In a previous study that used this metric for other targets,(Oh et al., 2014) projection strength values ≥10^−3.5^ (~0.0003) identified false axonal projections (primarily due to small segmentation artifacts) 14.5% of the time. At a higher threshold of 0.01, this false positive rate fell to ~0%.(Oh et al., 2014)^(extended^ ^Fig^ ^7)^ Thus, we used the 0.01 threshold for descriptive statistics. We also separately calculated the strength of contralateral and ipsilateral projections after repeating the search with left and right sided RF_motor_ targets, respectively. To identify CST projections for comparison, we repeated the above procedures with the medullary pyramids as the target structure (http://connectivity.brain-map.org/projection?searchMode=target&sourceDomain=315&primaryStructureOnly=true&tracers=10&isi=false&targetDomain=190&hemisphere=either&targetVolumeThreshold=0.0000).

To visualize the results in brain-space, we generated an anatomical surface model of the right hemi-brain without the olfactory bulb in GIFTI format from the 0.025 mm isotropic resolution CCFv3 atlas (using MATLAB R2017a and Connectome Workbench v.1.4.2). We then mapped the injection coordinates to the nearest surface vertex and labelled a 0.1 mm radius circle around those coordinates with the projection strength values for that injection. We also mapped the CCFv3 atlas labels for cortical regions of interest (ROIs) to the surface model, including the anterior cingulate area, medial prefrontal areas (orbital, prelimbic, infralimbic and frontal pole), secondary motor area, primary motor area and primary somatosensory area.

### Data analysis

Projection strength is not normally distributed and has a high frequency of zeros,(Oh et al., 2014) so nonparametric statistics were used for analysis. For each target (RF_motor_ & pyramids), projection strength was compared between cortical ROIs using Kruskal-Wallis ANOVA, followed by Mann-Whitney U pairwise comparisons with false discovery rate (FDR) correction(Benjamini and Hochberg, 1995) across ROIs. For each target, ipsilateral vs. contralateral projection strength was tested using Wilcoxon signed rank tests (paired by injection experiment). We also calculated a projection strength laterality index as: (ipsilateral – contralateral) / (ipsilateral + contralateral) * 100, which ranges from −100 (completely contralateral) to 100 (completely ipsilateral), where 0 indicates bilateral symmetry. Between targets, projection strength was compared across all injection experiments, for each cortical ROI, for ipsilateral & contralateral projections and for the laterality index. These between-target analyses used Wilcoxon signed rank tests (paired by injection experiment) with separate FDR correction across ROIs or lateralities. R statistical software(R Development Core Team, 2004) v3.6.0 was used for analysis. The significance threshold was p_FDR_<0.05.

### Exploratory trajectory comparison analysis

Presently, it is not possible to directly map the CRP trajectory from anterograde tracer experiments because injections that label CRP neurons likely also label neurons from other tracts. To address this issue, we performed an exploratory statistical analysis to identify brain voxels more likely to belong to CRP vs. CST projections. For each eligible injection experiment, we downloaded the 3-dimensional projection density image (at 0.05 mm isotropic resolution), which had been resampled to the CCFv3 atlas space.(Oh et al., 2014) Non-parametric permutation testing was then performed to test how strongly the projection density at each voxel was associated with the CRP vs. CST projection strength across injection experiments, while controlling for the total (whole-brain) projection strength of each experiment. This analysis used FSL randomise(Winkler et al., 2014) and threshold-free cluster enhancement(Smith and Nichols, 2009) with 5,000 permutations and a two-sided significance threshold of p_FDR_<0.05. Two-sided significance testing was performed by generating both contrast maps (RF_motor_ - pyramids and pyramids - RF_motor_), taking the minimum p-value in each voxel and multiplying it by two, then running FDR correction. For visualization, results were also projected onto the surface model of the right hemi-brain without the olfactory bulb.

## RESULTS

There were 607 eligible experiments injecting anterograde EGFP tracer into the right isocortex, among which 360 (59.3%) used male mice and 316 (52.1%) involved injections in the ROIs (Fig 1).

**Figure 1.**
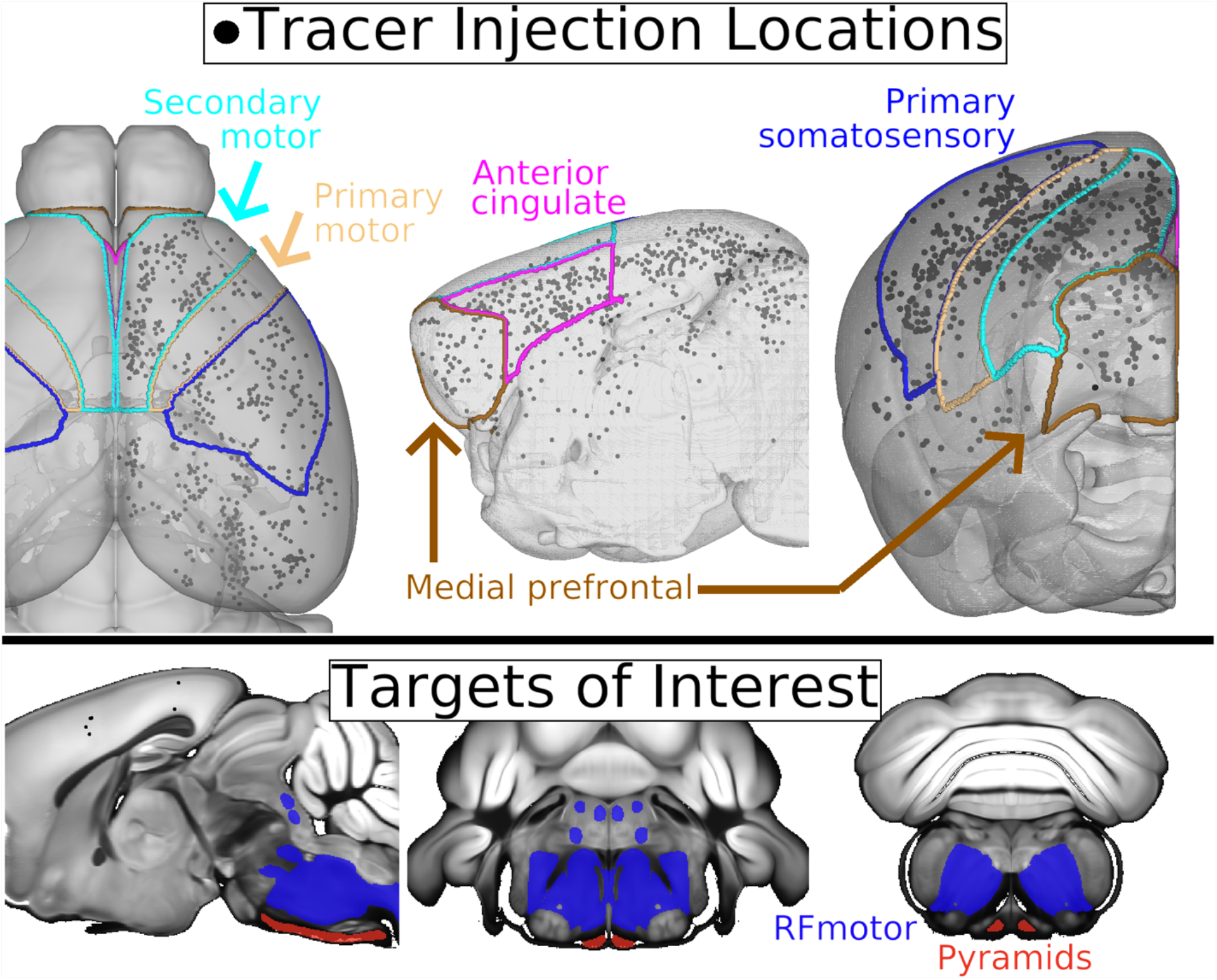
Locations of cortical anterograde tracer injections and brainstem targets of interest. **Top panel.** Each eligible injection experiment (N=607) is marked with a 0.1 mm diameter black sphere within the translucent brain surface rendering. Cortical regions of interest are outlined on the surface and annotated. In the middle and right images, the left hemi-brain and olfactory bulb have been removed from the surface rendering to better visualize the anterior cingulate and medial prefrontal areas. **Bottom panel.** From left to right, slices are x=0.4mm, y=-5.2mm and y=-7.3mm. RF_motor_, reticular formation motor nuclei involved with limb movement.

The proportion of cortical injections with RF_motor_ projection strengths ≥0.01 was 55.8% overall (Fig 2). Within the anterior cingulate, medial prefrontal, secondary motor, primary motor and primary somatosensory ROIs, this proportion was 78.8%, 73.9%, 80.8%, 87.5% and 64.7%, respectively (Fig 3A). The omnibus Kruskal-Wallis ANOVA revealed significant differences in RF_motor_ projection strength among ROIs (test statistic = 118.73, df = 5, p < 2.2 × 10^−16^). FDR-corrected pairwise comparisons found that each ROI had significantly greater RF_motor_ projection strength than other (non-ROI) cortical areas, and there were significant differences among ROIs (Table 2). The primary motor area had significantly greater RF_motor_ projection strength than all other ROIs except the secondary motor area, which had significantly greater projection strength than the medial prefrontal and primary somatosensory areas. The anterior cingulate area was not significantly different from the secondary motor, medial prefrontal or primary somatosensory areas. Compared with the pyramids target, projection strength to RF_motor_ was significantly greater overall and within all cortical ROIs except the primary somatosensory cortex, where the relative projection strengths were equivocal.

**Figure 2.**
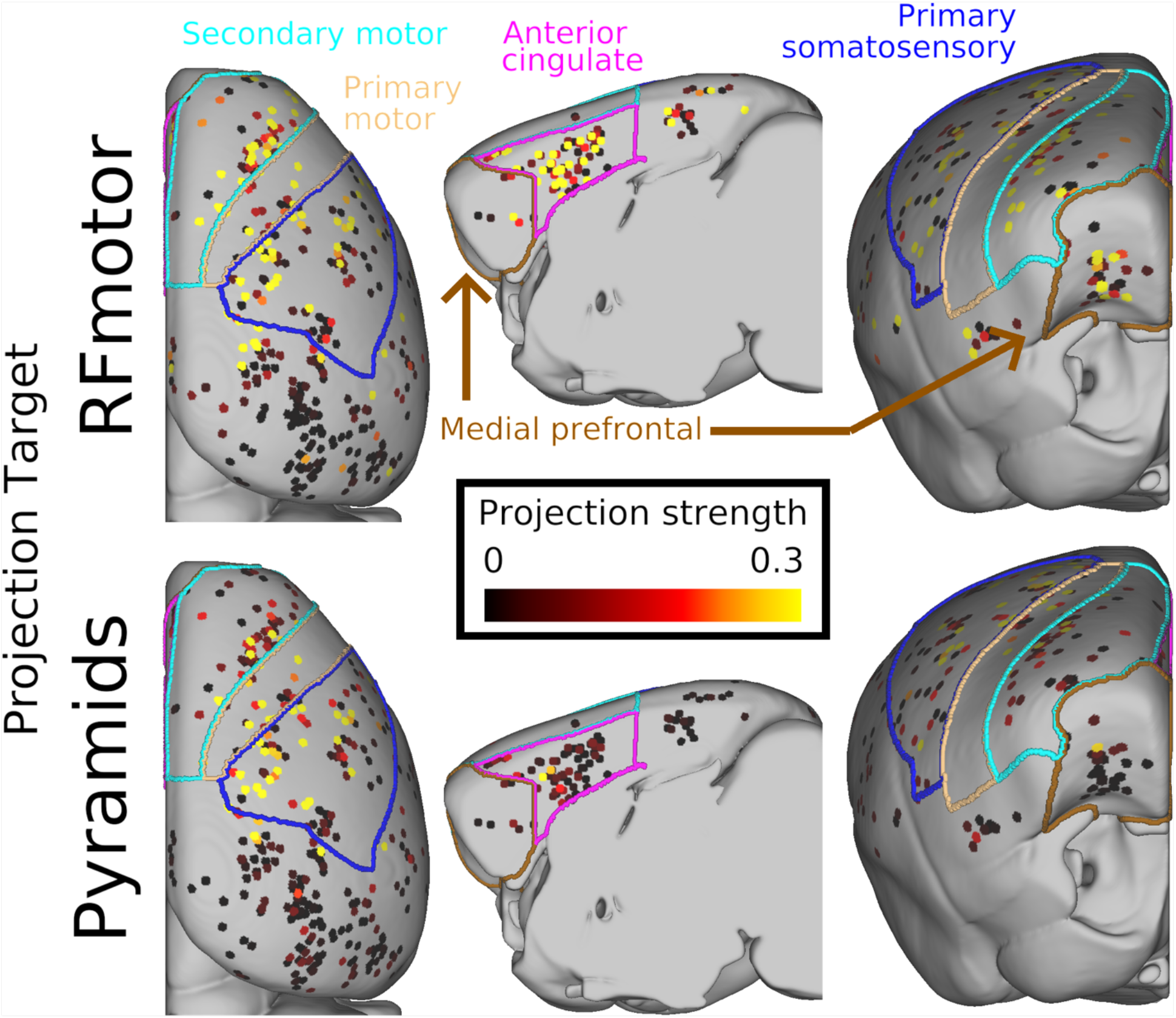
Anterograde tracer projections from cortical injection sites to the reticular formation motor nuclei (RFmotor) or medullary pyramids. Projection strength is the volume of tracer-labelled pixels in the target of interest (RF_motor_ or pyramids) divided by the injection site volume. Results from each eligible injection experiment (N=607) are mapped onto the nearest surface vertex with a 0.1 mm radius circle, color-coded with the projection strength from that cortical site to each brainstem target of interest. These results are mapped onto an opaque surface rendering of the right hemi-brain with the olfactory bulb removed for better visualization. Cortical regions of interest are outlined and annotated.

**Figure 3.**
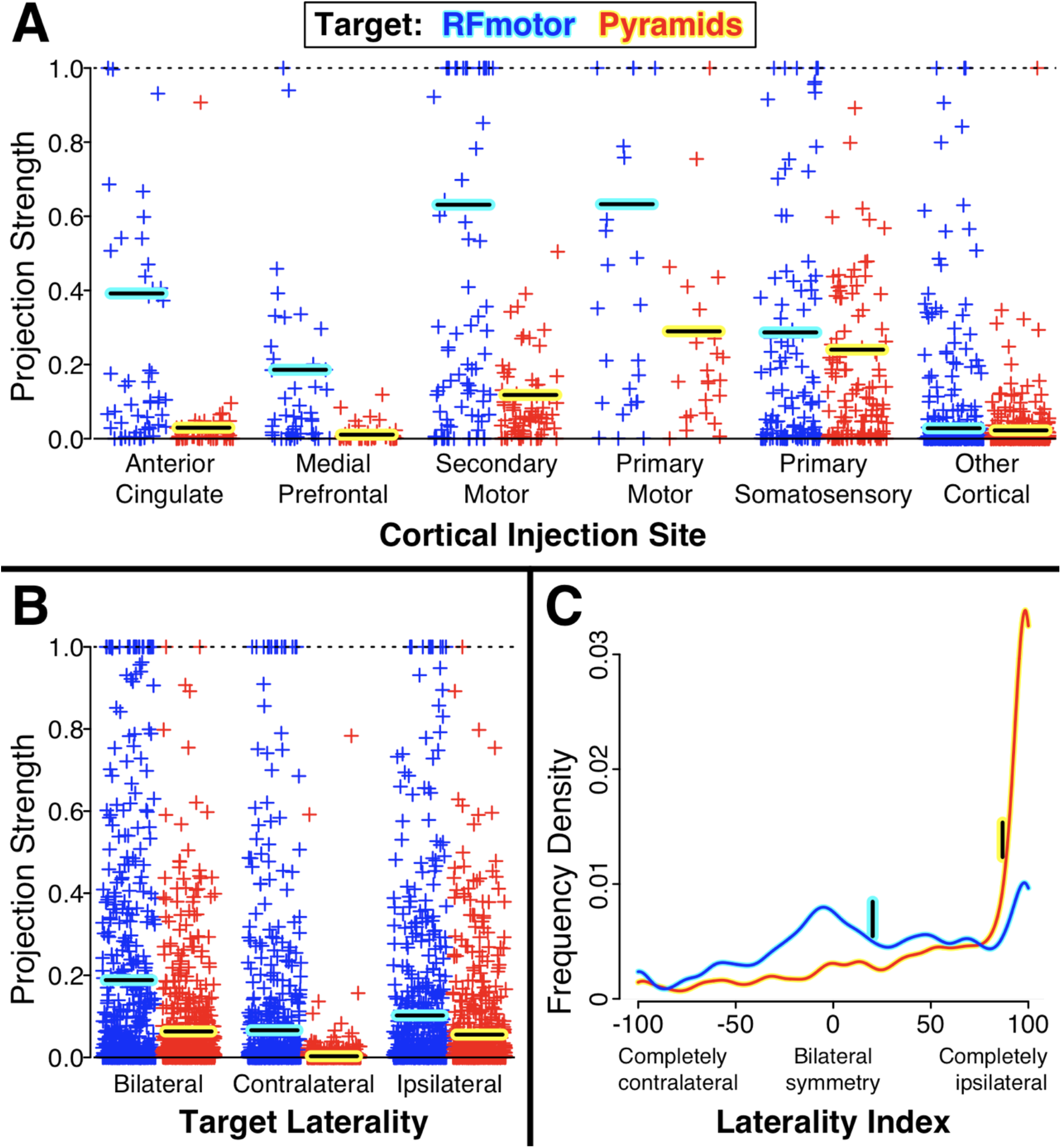
Projection strength to the brainstem targets by cortical injection site and target laterality. Projection strength is the volume of tracer-labelled pixels in the target divided by the injection site volume. **Panels A and B.** Each eligible injection experiment (N=607) is shown as a + symbol, with projection strength truncated at 1.0 for visualization. Horizontal bars indicate the 75^th^ percentile. **Panel C.** Distribution of tracer laterality across injection experiments for each target. Vertical bars indicate median laterality indices.

**Table 2.** Projection strength to the brainstem targets by cortical injection site and target laterality.

|  | Number of injections | Projection strength to **RF<sub>motor</sub>** |  |  | Projection strength to **Pyramids** |  |  | RF<sub>motor</sub> vs. Pyramids P<sub>FDR</sub>* |
| --- | --- | --- | --- | --- | --- | --- | --- | --- |
|  |  | Median (75%ile) | N (%) ≥0.01 |  | Median (75%ile) | N (%) ≥0.01 |  |  |
| All eligible experiments, bilateral targets | 607 | 0.02 (0.19) | 339 (55.8) |  | 0.01 (0.06) | 303 (49.9) |  | <0.001 |
| **By Cortical Injection Site** |  |  |  | P<sub>FDR</sub> vs. Other<sup>†</sup> |  |  | P<sub>FDR</sub> vs. Other |  |
| Anterior cingulate | 52 | 0.10 (0.39) | 41 (78.8) | <0.001<sup>b,c</sup> | 0.01 (0.03) | 22 (42.3) | 0.43<sup>c</sup> | <0.001 |
| Medial prefrontal | 46 | 0.05 (0.19) | 34 (73.9) | <0.001<sup>c</sup> | 0.00 (0.01) | 12 (26.1) | 0.56<sup>c</sup> | <0.001 |
| Secondary motor | 75 | 0.17 (0.63) | 60 (80.0) | <0.001<sup>a,b</sup> | 0.06 (0.12) | 60 (80.0) | <0.001<sup>b</sup> | <0.001 |
| Primary motor | 24 | 0.28 (0.63) | 21 (87.5) | <0.001<sup>a</sup> | 0.16 (0.29) | 22 (91.7) | <0.001<sup>a</sup> | 0.02 |
| Primary somatosensory | 119 | 0.05 (0.29) | 77 (64.7) | <0.001<sup>c</sup> | 0.08 (0.24) | 87 (73.1) | <0.001<sup>a,b</sup> | 0.18 |
| Other cortical | 291 | 0.00 (0.03) | 106 (36.4) | reference<sup>d</sup> | 0.00 (0.02) | 100 (34.4) | reference<sup>c</sup> | 0.001 |
| **By Target Laterality** |  |  |  | P vs. Ipsilateral |  |  | P vs. Ipsilateral |  |
| Contralateral | 607 | 0.01 (0.07) | 274 (45.1) | <0.001 | 0.00 (0.00) | 59 (9.7) | <0.001 | <0.001 |
| Ipsilateral | 607 | 0.01 (0.10) | 312 (51.4) | reference | 0.01 (0.06) | 278 (45.8) | reference | <0.001 |
| Laterality Index<sup>‡</sup> | 607 | 20.2 (68.8) | N/A | N/A | 86.8 (99.2) | N/A | N/A | <0.001 |

Projection strength = volume of tracer-labelled pixels in the target of (RF<sub>motor</sub> or pyramids) divided by injection site volume
\*P-values from Wilcoxon signed rank tests (paired by injection experiment) with false discovery rate (FDR) correction
<sup>†</sup>P-values from Mann-Whitney U tests with FDR correction
<sup>a,b,c,d</sup>Rows without a matching letter have pairwise comparisons with p<sub>FDR</sub><0.05
<sup>‡</sup>Ranges from -100 (completely contralateral) to 100 (completely ipsilateral), where 0 indicates bilateral symmetry

For the pyramids target, 49.9% of cortical injections had projection strengths ≥0.01. Within the anterior cingulate, medial prefrontal, secondary motor, primary motor and primary somatosensory ROIs, this proportion was 42.3%, 26.1%, 80.8%, 91.7% and 73.1%, respectively (Figs 2 & 3A, Table 2). The omnibus Kruskal-Wallis ANOVA revealed significant differences in pyramid projection strength among ROIs (test statistic = 148.10, df = 5, p < 2.2 × 10^−16^). FDR-corrected pairwise comparisons found that the motor and somatosensory areas had significantly greater pyramid projection strength than other (non-ROI) cortical areas, but the anterior cingulate and medial prefrontal areas did not (Table 2). The primary motor area had significantly greater pyramid projection strength than all other ROIs except the primary somatosensory area, which had significantly greater projection strength than the anterior cingulate and medial prefrontal areas. The secondary motor area was not significantly different from the primary somatosensory area and also had significantly greater projection strength than the anterior cingulate and medial prefrontal areas.

Ipsilateral projection strength was significantly greater than contralateral for both RF_motor_ and the pyramids (Fig 3B, Table 2). Compared with the pyramids target, projection strength to RF_motor_ was significantly greater for both the ipsilateral and contralateral projections. However, RF_motor_ projection strength was significantly less lateralized, with a median laterality index of 20.2, versus 86.8 for the pyramids (Fig 3C, Table 2).

In the exploratory CRP versus CST trajectory comparison analysis, we tested how strongly the projection density at each voxel was associated with RF_motor_ (CRP) versus pyramids (CST) projection strength, while controlling for total projection density. A large cluster of voxels with significantly greater RF_motor_ association spanned all right cortical ROIs and was anterior to the main cluster of greater pyramids association, which included the medial parts of the primary motor and primary somatosensory areas (Fig 4 row 1). In the subcortex, the cluster of greater RF_motor_ association followed a trajectory towards the brainstem through the internal capsule that was anterior, ventral and medial to the cluster of greater pyramids association (Fig 4 rows 2-6). In the ventral diencephalon and rostral midbrain, the two trajectories crossed and the cluster of greater RF_motor_ association became dorsal to the cluster of greater pyramids association. The cluster of greater RF_motor_ association then expanded to fill most of the dorsal pons and medulla bilaterally, with multiple areas of apparent decussation, especially in the pons. Meanwhile, the cluster of greater pyramids association followed the compact trajectory of the pyramids in the ventral brainstem and remained primarily ipsilateral before most of the cluster began decussating in the most caudal slices of the medulla.

**Figure 4.**
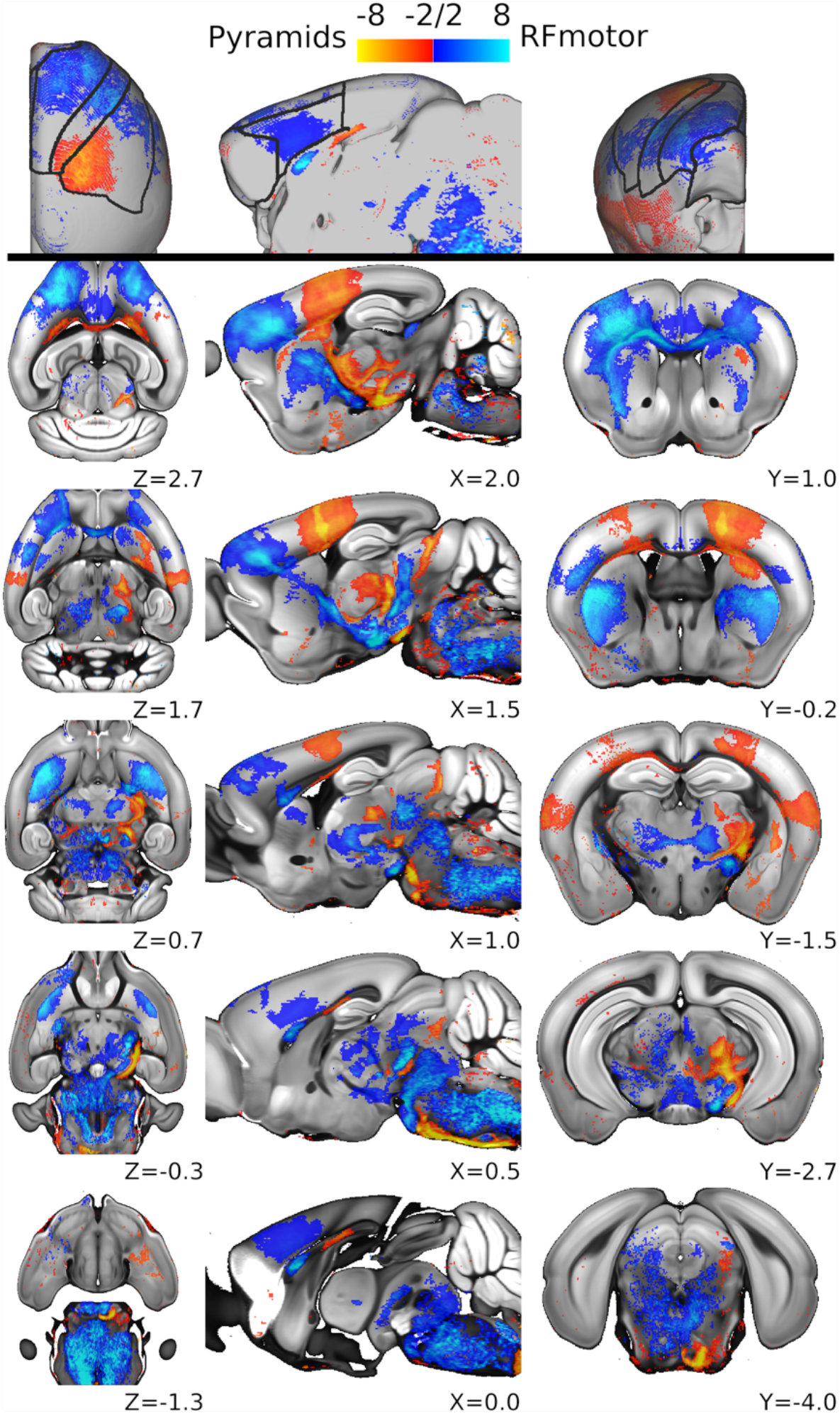
Exploratory trajectory analysis of the corticoreticular pathway (CRP) versus the corticospinal tract (CST). Mapped results are T statistics from a general linear model of projection density at each voxel, testing its association with the projection strength to the reticular formation motor nuclei (RF_motor_) versus the medullary pyramids, while controlling for whole-brain projection strength (N=607). These results are from non-parametric permutation testing using threshold-free cluster enhancement, and are thresholded at a two-sided, false discovery rate corrected p<0.05. Blue T statistics indicate a significantly greater association with RF_motor_ (CRP) projection strength, while red/yellow T statistics indicate a significantly greater association with pyramids (CST) projection strength. The analysis was done in volume space (rows 2-6) and these volumetric images are in neurologic orientation. For visualization, results were also projected onto the surface model of the right hemibrain without the olfactory bulb (row 1). This surface visualization includes black outlines of the cortical regions of interest shown in other figures.

## DISCUSSION

This study used cortex-wide anterograde axonal tracing data to map the mouse CRP, in comparison with the CST, with an emphasis on cortical inputs and laterality. As expected, the motor cortex provided strong CRP and CST projections. The CRP received its strongest inputs from the primary and secondary motor areas while the CST received its strongest inputs from the primary motor and primary somatosensory areas. Unlike the CST, the CRP also had strong projections originating from the anterior cingulate and medial prefrontal areas. Both tracts had significantly stronger ipsilateral vs. contralateral projections (above the level of the pyramidal decussation), but the CRP was less lateralized, with a greater proportion of bilateral projections. The CRP had greater projection strength than the CST for all ROIs except the primary somatosensory cortex, and for both ipsilateral and contralateral projections.

Overall, these results indicate that the CRP is a widely distributed, bilaterally projecting tract in the mouse with more diverse cortical inputs than the CST and thus greater likelihood of partial sparing after brain injury. This is consistent with a recent study that found loss of ipsilesional CST projection strength and upregulation of the ipsilesional CRP, which was correlated with motor recovery after frontoparietal lesions in primates.(Darling et al., 2018) It is also consistent with another recent study showing residual capacity for ipsilesional CRP upregulation following intracerebral hemorrhage in rats.(Ishida et al., 2019) In addition, the bilateral projection strength of the CRP is reinforced by studies in mice(Takase et al., 2017) and humans(Jang et al., 2013) reporting upregulation of the contralateral CRP after stroke, which was correlated with motor recovery.

The current findings appear to diminish long-standing uncertainty from classical tract tracer studies about whether there are CRP projections emanating from the anterior cingulate and medial prefrontal cortices.(Berrevoets and Kuypers, 1975; Keizer et al., 1984; Keizer et al., 1989; Newman et al., 1989; Rho et al., 1997; Rossi and Brodal, 1956) From our large, cortex-wide analysis of next-generation anterograde tracer experiments, it now seems clear that each of these projections is relatively strong, at least in the mouse. Many prior animal studies of the CRP have focused solely on its strongest origins in the motor cortex.(Asboth et al., 2018; Darling et al., 2018; Fisher et al., 2021; Fregosi et al., 2017; He and Wu, 1985; Ishida et al., 2019; Jinnai, 1984; Kably and Drew, 1998; Kably and Drew, 1998; Lamas et al., 1994; Matsuyama et al., 2004; Pilyavsky, 1975; Takase et al., 2017) Thus, future studies are needed to determine the extent to which these additional portions of the CRP might be capable of mediating motor recovery after a brain injury.

Our findings also raise pressing questions about the degree to which CRP projections from the anterior cingulate and medial prefrontal cortices may have been preserved in humans. If these projections have been evolutionarily preserved, prior diffusion tractography studies have vastly underestimated the extent of the CRP. Several of these studies have restricted the analysis to projections involving either the precentral gyrus(Lindenberg et al., 2010; Schulz et al., 2017; Zheng and Schlaug, 2015) or secondary motor cortex,(Jang and Seo, 2015; Yeo et al., 2012; Yeo et al., 2020) possibly because initial attempts to map the human CRP without limiting the cortical search window have not found anterior cingulate or medial prefrontal origins.(Soulard et al., 2020; Yeo et al., 2014) However, these studies performed tractography with low-resolution diffusion-weighted imaging and the simple diffusion tensor model, which is unable to resolve multiple fiber populations within a voxel and thus highly prone to false negatives (and false positives).(Calamante, 2019) Using a slightly more sophisticated model, Jang and Seo(Jang and Seo, 2014) identified CRP streamlines originating from the dorsal prefrontal cortex, but did not search the anterior cingulate cortex or as far anterior as the medial prefrontal cortex. Thus, it is still possible that CRP bundles originating from the anterior cingulate and medial prefrontal cortices may have persisted in humans. Assessing this possibility with higher resolution diffusion-weighted imaging and next generation modeling methods should be a priority.

Given that some prior studies have only studied CRP projections originating from the secondary motor area,(Jang and Seo, 2015; Yeo et al., 2012; Yeo et al., 2020) another important finding was that the primary and secondary motor areas had similar CRP projection strength. Prior tract tracer and invasive neurophysiology studies in animals (including primates) have also consistently found evidence of robust CRP projections originating from the primary motor cortex, including many collateral branches of CST axons.(Asboth et al., 2018; Fisher et al., 2012; Fisher et al., 2021; Fregosi et al., 2017; He and Wu, 1985; Ishida et al., 2019; Jinnai, 1984; Kably and Drew, 1998; Kably and Drew, 1998; Lamas et al., 1994; Matsuyama et al., 2004) Human CRP tractography studies have identified these primary motor cortex projections too.(Jang and Seo, 2014; Lindenberg et al., 2010; Schulz et al., 2017; Zheng and Schlaug, 2015) This indicates that primary motor cortex stimulation or recording likely does not specifically target the CST as often presumed.^e.g.^(Barthélemy et al., 2011; Capaday et al., 1999; Chieffo et al., 2016; Jayaram et al., 2012) It also suggests that projections from the primary motor area should not necessarily be omitted during CRP tractography.

Our broad quantification of CRP laterality (Fig 3B & 3C; Table 2) is consistent with the more focal results from prior studies.(Fisher et al., 2021; Kably and Drew, 1998; Rho et al., 1997) For example, Kably and Drew(Kably and Drew, 1998) used microstimulation to measure the laterality of 157 CRP neurons originating from primary or secondary motor areas in the cat and found that 49% were ipsilateral, 35% were bilateral and 16% were contralateral. Using retrograde axonal tracing in the cat, Rho et al(Rho et al., 1997) found that the percentage of ipsilateral CRP projections varied from 46.0% to 72.9% across different RF_motor_ nuclei. In a recent microstimulation study among primates, Fisher et al(Fisher et al., 2021) found bilateral CRP projections to 20/36 (56%) RF_motor_ neurons with inputs from the primary motor cortex and 30/36 (83%) RF_motor_ neurons with inputs from the secondary motor cortex. The current analysis extended these findings by showing the distribution of CRP laterality across cortical injection sites (with high proportions of both ipsilateral and bilateral projection strength) and by quantitatively confirming that the CRP is significantly less lateralized than the CST.

In our exploratory voxel-wise analysis testing projection density associations with RF_motor_ (CRP) versus pyramids (CST) projection strength, the main statistically significant clusters followed paths consistent with plausible CRP trajectories and known CST trajectories (Fig 4). If these statistical results for the CRP are consistent with actual axonal trajectories, it would indicate that the CRP runs anterior, medial and ventral to the CST in the subcortical white matter then moves dorsal to the CST in the ventral diencephalon and rostral midbrain. Interestingly, this is consistent with results from preliminary subcortical CRP mapping with diffusion tractography in humans.(Jang and Seo, 2015) Our results also suggest that the CRP projects bilaterally throughout the dorsal pons and medulla, with multiple decussation points, especially in the pons.

However, other significant association clusters did not match plausible CRP or CST trajectories. The majority of these were in cerebral commissures and contralateral cerebral gray matter regions that were homotopic with ipsilateral clusters and did not project to the brainstem. Thus, we suspect that these clusters were due to confounding from commissural projections that happened to originate near CRP and CST projection neurons, or as collateral branches of CRP or CST axons. When interpreting this analysis, another key consideration is that it was a contrast between RF_motor_ and pyramids projection associations and thus it could not find areas of CRP and CST overlap.

### Limitations

An important limitation to this study is that there was incomplete coverage of some cortical subregions, despite the large number of anterograde injection experiments. There was also sparser coverage of the primary motor cortex compared with other ROIs. Thus, next-generation retrograde tracer experiments may be able to provide more granular CRP mapping in the future. Another possible issue is that we were only able to include injections in the right isocortex, but there have not been strong indications of interhemispheric CRP differences in prior studies. The projection strength metric used in this analysis does not differentiate tract terminations from continuations, so it could be falsely elevated by sparse fibers passing through RF_motor_ or the pyramids without synapsing. Conversely, using transgenic mice and injections at various cortical depths may have falsely lowered projection strength for some experiments, resulting in random measurement error. However, this is unlikely to have caused any systematic error (bias) in the analyses because the CRP and CST results were obtained from the same injections and there is no reason to suspect systematic differences between ROIs.

## Conclusions

The mouse CRP bilaterally converged on RF_motor_ from large portions of the cortex, including the primary & secondary motor areas, anterior cingulate, primary somatosensory area and medial prefrontal area, in order of decreasing projection strength. Compared with the CST, the CRP was less lateralized and had stronger projections from all these cortical regions except the primary somatosensory cortex. In the subcortex, the CRP appeared to descend anterior, ventral and medial to the CST before moving dorsal to the CST near the rostral midbrain and projecting bilaterally throughout the dorsal pons and medulla. These findings theoretically increase the likelihood of partial CRP sparing after brain injury and reinforce the conceptual basis for contralesional CRP compensation after unilateral damage. This foundational information can be used to guide future CRP tractography and projection-specific manipulations. The current study also highlights the value of robust connectomic data generation and sharing(Oh et al., 2014) for enabling ancillary analyses to accelerate scientific progress.

## AUTHOR CONTRIBUTIONS

Conceptualization, Resources, Software, Formal analysis, Visualization, Writing - original draft: PB. Methodology, Writing - review and editing: PB, OOA, YL.

## DATA ACCESSIBILITY STATEMENT

The data used for this analysis are available from the Allen Mouse Brain Connectivity Atlas (© 2011 Allen Institute for Brain Science http://connectivity.brain-map.org/).

